# Thermal adaptation and fatty acid profiles of bone marrow and muscles in mammals: implications of a study of caribou (*Rangifer tarandus caribou*)

**DOI:** 10.1101/2022.05.04.490689

**Authors:** Eugène Morin, Päivi Soppela, P. Yvan Chouinard

**Affiliations:** Trent University, Department of Anthropology, DNA Bldg Block C, 2140 East Bank Drive, Peterborough, Ontario, Canada K9J 7B8; Université de Bordeaux, PACEA, Bat. B18, Allée Geoffroy St-Hilaire CS 50023, 33615 Pessac Cedex, France; Arctic Centre, University of Lapland, Pohjoisranta 4, P.O. Box 122, 96101 Rovaniemi, Finland; Université Laval, Département des sciences animales, Pavillon Paul-Comtois, Québec, Canada G1V 0A6

**Keywords:** thermal adaptation, fatty acids, lipids, bone marrow, meat, muscles, intramuscular fat

## Abstract

Mammals have evolved several physiological mechanisms to cope with changes in ambient temperature. Particularly critical among them is the process of keeping cells in a fluid phase to prevent metabolic dysfunction. In this paper, we examine variation in the fatty acid composition of bone marrow and muscle tissues in the cold-adapted caribou (*Rangifer tarandus caribou*) to determine whether there are systematic differences in fatty acid profiles between anatomical regions that could potentially be explained by thermal adaptation. Our results indicate that the bone marrow and muscle tissues from the appendicular skeleton are more unsaturated than the same tissues in the axial skeleton, a finding that is consistent with physiological adaptation of the appendicular regions to thermal challenges. Because mechanisms of thermal adaptation appear to be widely shared among terrestrial mammals, we suggest that the same patterns may prevail in other species, possibly including humans.

## Introduction

How mammals, including humans, adapt to changes in ambient temperature has long been a focus of intensive research in biology (Schmidt-Nielsen 1946; Irving and Krog 1955; Irving et al. 1957; Meng et al. 1969; Soppela et al. 1986). A critical challenge that all mammals must face is to maintain their high internal body temperature by conserving heat in cold weather and dissipating heat in hot weather and/or when the animal is conducting long, vigorous activity (Blix 2005). At the scale of individual cells, the problem concerns how the physical properties of the membrane and its composition can be modified in order to dynamically maintain a fluid (liquid-crystalline) phase under challenging thermal conditions (Hochachka and Somero 2002). At low temperatures, this means keeping the cell away from a gel phase, whereas at high temperature it implies avoiding the development of inverted hexagonal phase structure and membrane fusion (Hazel 1995). Avoiding these changes in phase or structure is crucial because they can have deleterious effects on cell function (Stillwell 2016), and at a larger scale, may result in stiff or loose tissues, with potentially adverse effects on locomotion, food procurement and the ability of an animal to respond swiftly in contexts of predation. In conditions of low ambient temperature, a common pattern seen in cells is desaturation, which consists in increased proportion of unsaturated fatty acids (FA) at the expense of saturated FA. At high ambient temperatures, these changes are commonly reversed (Hochachka and Somero 2002; Denlinger 2010; Stillwell 2016, Pond 2017). In this paper, we examine how the FA composition of skeletal marrow from different parts of the body varies—likely in response to thermal challenges (Meng et al. 1969)—in a species of cold climate, the caribou (*Rangifer tarandus caribou*). The FA composition in bone marrow is also compared with that of skeletal muscles to determine whether this tissue is similarly affected by exposure to ambient temperature.

Bone marrow adipocytes (BMA) are metabolically active cells that are currently intensively studied because they act as an energy reservoir, secrete important proteins (e.g., adiponectin, leptin) and influence local marrow processes, osteogenesis and systemic metabolism, among other functions (Li et al. 2018; Hawkes and Mostoufi-Moab 2019; Weldenegodguad et al. 2021). In humans, the formation of BMA occurs at or shortly prior to birth with these cells gradually replacing hematopoietic tissue. This process follows a well established distal-to-proximal sequence: BMA first form in the terminal phalanges, then develop in the long bones, and later expand into the axial skeleton (Tavassoli and Yoffey 1985). Within individual long bones, BMA first form at mid-shaft, then occur in the distal metaphysis, and ultimately invade the proximal metaphysis (Kricun 1985; Moore and Lawson 1990). Importantly, these developmental trends are not unique to humans and have been observed in many mammal species, including rodents, ungulates, and carnivores (e.g., Goodman 1952; Day 1977).

Recent studies have uncovered important molecular, functional and morphological differences between subtypes of BMA. In the metaphyseal regions of the long bones—loci generally associated with active hematopoiesis—BMA represent only approximately 45% of the cellular component in adults, which contrasts with their abundance in long bone cavities where they can constitute as much as 90% of the cellular tissue (Snyder et al. 1975; Lecka-Czernik et al. 2018). The BMA found in hematopoietic regions also tend to be of smaller size, to be more dispersed, to develop later and to be more easily mobilized than the BMA found in the shaft cavities of the same bones. Moreover, BMA in metaphyseal and diaphyseal regions have been shown to differ in terms of patterns of regulation and gene expression (Scheller et al. 2015; Craft et al. 2018). As a result of these differences, the adipocytes from the metaphyseal regions have been termed regulated BMA (rBMA, or “red marrow” in the older literature), whereas those found in the diaphyses or shafts of long bones are called constitutive BMA (cBMA, or “yellow marrow” in early publications) (Scheller et al. 2015).

Research on terrestrial mammals has shown that the FA composition of BMA is influenced by several factors, including the anatomical locus under investigation as well as the age, diet, health, and gender of the individual (Tavassoli and Yoffey 1985; Soppela and Nieminen 2001, Huovinen et al. 2015, Steiner-Bogdaszewska et al. 2022). Because the FA composition of BMA appears to be influenced by tissue temperature, and indirectly by ambient temperature (Meng et al. 1969; Turner 1979; Pond et al. 1993; Käkelä and Hyvärinen 1996, Soppela and Nieminen 2001), comparing patterns in the axial skeleton with that from the more heterothermic limbs may yield important insights on the interactions between cell function and thermal challenges. For instance, cBMA are known, to increase in unsaturation towards the extremities, and are thus kept fluider, in order to prevent stiffness at low temperatures (Meng et al. 1969; Turner 1979; Pond et al. 1993; Käkelä and Hyvärinen 1996; Soppela and Nieminen 2001). Whether this pattern also applies to the adjacent muscles and to cells in the metaphyseal regions is poorly known. This issue is important because, unlike cBMA, intramuscular fat in lean animals has a high content of structural lipids such as phospholipids and cholesterol (Leat and Cox 1980; Lawrie and Ledward 2006; Wood et al. 2008), an observation that likely extends to the marrow in the metaphyseal regions given its hematopoietic functions. How the FA composition of muscle and bone marrow tissues varies within an animal and how these tissues are influenced by ambient temperature is understudied (but see Irving and Krog 1955; Hammel et al. 1962; Johnsen et al. 1985). The present study addresses this problem by investigating changes in the FA composition of these tissues across a large number of different anatomical sites in caribou. We also explore the implications of these variations for our understanding of the thermal adaptation of mammals.

## Materials and Methods

### Sample collection

Whereas many previous studies of FA composition in wild mammals have investigated a single category of tissue (e.g., adipose tissue, bone marrow or a specific skeletal muscle) sampled across a large number (e.g., >10) of individuals at a small number of anatomical sites (e.g., 1–5 sites), here we used a different approach and focused on intra-individual variation. For this study, FA variation was examined at a large number of anatomical sites (*n*=56 per individual) in caribou, with special attention being paid to a wide range of soft tissues, including backfat, skin, skeletal muscle, lungs and trachea, and bone marrow. This sampling strategy allows for a detailed picture of FA profile variation within a cold-adapted mammal.

For this analysis, two already eviscerated female (≥6 year-old) caribou (*Rangifer tarandus caribou*) were sampled. Both free-ranging caribou were killed by hunters around February 7–10, 2007 in the Robert-Bourassa Reservoir (53°45’N, 77°00′W; female A) and the Caniapiscau region (53°00′N, 68°30′W; female B) in central Québec where average temperature is around -23°C in January and 13°C in July (Schefferville airport weather station). Carcass weights (excluding organs, visceras, brain and antlers) were 64 and 61 kg, respectively. The two animals were, based on visual inspection, apparently in good condition and showed no signs of pathology or being starved. Whether the animals were pregnant or not is unknown. As part of the meat aging process, carcasses were kept in a refrigeration facility (at ca 4°C) for about two weeks prior to processing by a commercial butcher. Fatty acid profiles were derived for a total of 112 samples (56 per caribou) collected from the skin, lungs and trachea, various muscles, and the bone marrow from most classes of skeletal elements (in the case of long bones, samples were taken from the metaphyseal and shaft portions of the bones). Figure 1 shows the anatomical location of the samples.

**Fig 1.**
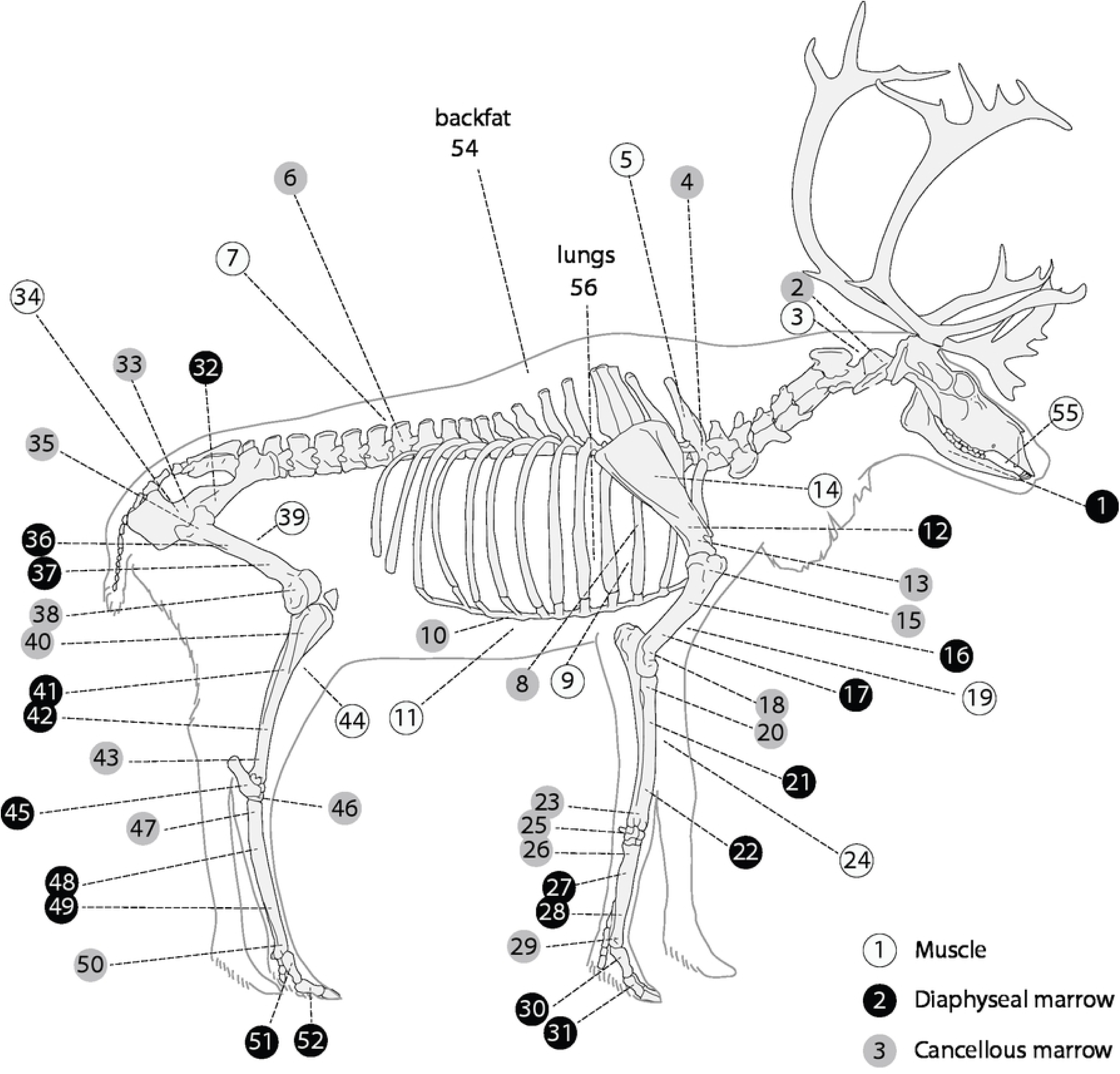
Anatomical sites sampled for this study. The numbers correspond to tissues listed in Table 1.

### FA extraction and analysis

How the marrow was obtained requires additional description. Because the FA analysis was performed with the aim of shedding light on human foraging decisions in prehistoric contexts—a topic that will be the focus of another publication—the marrow from the shaft cavity (diaphyseal marrow) of the long bones (humerus, radio-ulna, femur, tibia and metapodials) was extracted after breaking the bones using a pebble and a stone anvil as documented in a wide range of subsistence-based societies (Morin 2020). After the long bone shaft cavity was breached, a few grams of the exposed diaphyseal marrow was cut using a knife and then frozen in a plastic bag in a commercial freezer prior to FA analysis. One sample each was taken from the proximal and distal ends of the marrow plug. The cancellous marrow samples (marrow in the axial skeleton and girdles, carpals, tarsals, and metaphyseal regions of long bones; Fig 1) were obtained by crushing the specimen using the same stone-and-anvil method. This crushing yielded a product similar to bone meal or bone paste. This means that while the marrow extracted from the shaft cavity of the long bones was largely fat-like (or fatty) tissue and free of bone fragments, the crushed cancellous bone samples consisted of marrow-rich bone meal. As the bone tissue itself—that is excluding the soft tissue found in the trabeculae—contains very little fat (Higgs et al. 2011), endogenous bone fat is not expected to impact our results. The muscle samples consisted of small remnants of meat adhering to the bone, which were cut off after the main muscle masses had been removed by the butcher.

**Table 1.** Crude fat and fatty acid content of various body tissues in caribou. The values are averages for the two animals.

| Anatomical site | Tissue (Reference #) | Crude fat<sup>1</sup><br>(%, wet basis) | Total fatty acids<br>(%, crude fat basis) | Total fatty acids<sup>1</sup><br>(%, wet basis) |
| --- | --- | --- | --- | --- |
| Mandible | Marrow (1) | 60.1 | 78.0 | 46.8 |
| Cervical vertebrae | Cancellous marrow (2) | 28.0 | 70.9 | 19.9 |
|  | Meat (3) | 5.5 | 69.8 | 3.9 |
| Thoracic vertebrae | Cancellous marrow (4) | 21.2 | 79.2 | 16.8 |
|  | Meat (5) | 5.0 | 65.5 | 3.3 |
| Lumbar vertebrae | Cancellous marrow (6) | 23.2 | 76.5 | 17.8 |
|  | Meat (7) | 7.3 | 64.3 | 4.7 |
| Rib | Cancellous marrow (8) | 17.2 | 75.9 | 13.0 |
|  | Meat (9) | 9.7 | 84.1 | 7.5 |
| Sternum | Cancellous marrow (10) | 32.2 | 68.0 | 21.5 |
|  | Meat (11) | 26.6 | 62.4 | 17.2 |
| Scapula | Marrow (12) | 51.2 | 79.8 | 40.7 |
|  | Cancellous marrow (13) | 12.1 | 77.2 | 9.3 |
|  | Meat (14) | 4.8 | 54.3 | 2.6 |
| Humerus | Prox. metaphysis cancellous (15) | 38.0 | 67.0 | 25.4 |
|  | Prox. shaft marrow (16) | 84.6 | 72.6 | 61.3 |
|  | Distal shaft marrow (17) | 75.1 | 75.5 | 56.8 |
|  | Distal metaphysis cancellous (18) | 19.1 | 67.9 | 13.1 |
|  | Meat (19) | 6.3 | 42.1 | 2.7 |
| Radio-ulna | Prox. metaphysis cancellous (20) | 14.5 | 70.4 | 10.6 |
|  | Prox. shaft marrow (21) | 75.4 | 85.2 | 64.0 |
|  | Distal shaft marrow (22) | 76.9 | 84.9 | 65.1 |
|  | Distal metaphysis cancellous (23) | 15.6 | 71.4 | 11.2 |
|  | Meat (24) | 11.3 | 17.7 | 2.0 |
| Carpals | Cancellous marrow (25) | 10.4 | 69.6 | 7.1 |
| Metacarpal | Prox. metaphysis cancellous (26) | 7.5 | 69.1 | 5.1 |
|  | Prox. shaft marrow (27) | 70.2 | 81.6 | 57.2 |
|  | Distal shaft marrow (28) | 64.5 | 77.6 | 50.1 |
|  | Distal metaphysis cancellous (29) | 9.5 | 74.6 | 7.2 |
| Anterior phalanx 1 | Marrow (30) | 75.1 | 78.8 | 59.2 |
| Anterior phalanx 2 | Marrow (31) | 74.3 | 77.6 | 57.7 |
| Pelvis | Marrow (32) | 68.9 | 69.7 | 47.6 |
|  | Cancellous marrow (33) | 24.1 | 80.0 | 19.4 |
|  | Meat (34) | 4.2 | 56.1 | 2.3 |
| Femur | Prox. metaphysis cancellous (35) | 32.7 | 77.4 | 25.3 |
|  | Prox. shaft marrow (36) | 75.3 | 81.2 | 61.2 |
|  | Distal shaft marrow (37) | 74.3 | 85.1 | 63.1 |
|  | Distal metaphysis cancellous (38) | 30.5 | 81.6 | 24.9 |
|  | Meat (39) | 10.2 | 62.3 | 6.4 |
| Tibia | Prox. metaphysis cancellous (40) | 30.6 | 77.3 | 23.1 |
|  | Prox. shaft marrow (41) | 78.9 | 85.7 | 67.6 |
|  | Distal shaft marrow (42) | 72.2 | 78.7 | 56.8 |
|  | Distal metaphysis cancellous | 20.7 | 79.3 | 16.4 |
|  | Meat (43) | 7.0 | 78.0 | 5.4 |
| Calcaneus | Marrow (44) | 55.7 | 85.9 | 46.5 |
| Tarsals | Cancellous marrow (45) | 7.7 | 73.8 | 5.7 |
| Metatarsal | Prox. metaphysis cancellous (46) | 12.7 | 72.6 | 9.2 |
|  | Prox. shaft marrow (47) | 79.0 | 74.9 | 59.1 |
|  | Distal shaft marrow (48) | 72.3 | 79.5 | 57.1 |
|  | Distal metaphysis cancellous (49) | 15.9 | 77.1 | 12.2 |
| Posterior phalanx 1 | Marrow (51) | 71.5 | 80.2 | 57.3 |
| Posterior phalanx 2 | Marrow (52) | 63.5 | 83.2 | 52.8 |
| Miscellaneous | Skin (53) | 3.5 | 21.3 | 0.8 |
|  | Backfat (54) | 89.8 | 74.7 | 66.8 |
|  | Tongue (55) | 33.1 | 71.3 | 23.7 |
|  | Lungs and windpipe (56) | 10.0 | 80.2 | 7.7 |
| Mean and st. dev. | Cancellous marrow<sup>1-2</sup> | 20.1 ±9.1 | 74.5 ±4.5 | 15.0 ±6.7 |
|  | axial skeleton only<sup>1</sup> | 19.5 ±8.3 | 74.6 ±4.3 | 14.5 ±5.9 |
|  | metaphyseal regions only<sup>1</sup> | 20.6 ±10.0 | 73.8 ±4.8 | 15.3 ±7.5 |
|  | Diaphyseal marrow<sup>3</sup> | 74.9 ±5.0 | 80.2 ±4.5 | 60.0 ±4.7 |
|  | Meat | 10.9 ±9.2 | 60.7 ±17.4 | 6.8 ±6.7 |

<sup>1</sup>The low values for crude fat and Total fat weight (%, wet basis) for the cancellous marrow are, in part, due to the presence of bone fragments in the samples.
<sup>2</sup>Cancellous marrow includes the following: vertebrae, ribs, sternum, scapulae, pelvis, metaphyseal regions of long bones, carpals, tarsals. The axial skeleton includes all of these bones to the exclusion of the long bones.
<sup>3</sup>Diaphyseal marrow includes all of the shaft portions of the long bones. Although not “true” long bones, metapodials are treated here as long bones due to their large marrow cavity.

Prior to the FA analysis, all tissues were manually triturated with a scalpel to obtain a homogenized product. The crude fat—which includes all types of lipids (triacylglycerols, phospholipids and cholesterol esters)—from the muscle and marrow samples was extracted using the method presented in Folch et al. (1957), as modified by Dryer et al. (1970) who recommended adding methanol in two separate aliquots in the initial steps of the procedure. Total FA (different lipid classes were not separated) in extracted lipids were then transesterified according to the method described by Chouinard et al. (1997) using 100 μl of 0.5 *M* Na in methanol per ca 10 mg fat in 1 mL of hexane. Determination of the fatty acid profile was carried out with a gas chromatograph (HP 5890A Series II, Hewlett Packard, Palo Alto, CA) equipped with a 100-m CP-Sil 88 capillary column (Chrompack, Middelburg, the Netherlands) and a flame ionization detector, as described by Faucitano et al. (2008). The melting point of fat extracted from each sample was estimated as the weighted sum of the melting point of individual FA, as described by Toral et al. (2013). The comparisons that we performed include an examination of different indices, including the Δ^9^ desaturase index, the percentage of polyunsaturated FA (PUFA), the percentage of short chain saturated FA (Fig 3c) and the n-6/n-3 ratio. How these were calculated is presented in the accompanying figures and tables. For comparative purposes, we derived melting points from the caribou FA profiles published by Meng et al. (1969). Note that there are systematic differences between the two datasets likely because their earlier analyses could not identify certain categories of FA that can now be routinely identified, thanks to progress in gas chromatograph technology. The results that we present below begin with the limbs, as the FA composition of these body parts has previously been shown to be influenced by exposure to ambient temperature. For comparison purposes, we calculated a number of percentages and ratios. The summed percentage of PUFA (%PUFA) includes the following FA: *cis*-9,12 18:2; *cis*-9,12,15 18:3; *cis*-6,9,12,15 18:4; *cis*-11,14 20:2; *cis*-8,11,14 20:3; *cis*-5,8,11,14 20:4; *cis*-5,8,11,14,17 20:5; *cis*-7,10,13,16 22:4; *cis*-4,7,10,13,16 22:5; *cis*-7,10,13,16,19 22:5; *cis*-4,7,10,13,16,19 22:6. The percentage of short chain saturated FA (%short chain saturated FA) focuses on the summed presence of 14:0 + 15:0. The following equation was used to derive the Δ^9^ desaturase index: (*cis*-9 14:1 + *cis*-9 16:1 + *cis*-9 18:1)/(14:0 + *cis*-9 14:1 + 16:0 + *cis*-9 16:1 + 18:0 + *cis*-9 18:1). This index was selected because it includes FA that are likely to have a significant impact on melting point. The n-6/n-3 ratio was calculated using this formula: n-6 PUFA/n-3 PUFA.

### Statistical analysis

Differences in means between diaphyseal and other bone regions were assessed using unpaired two-tailed t-tests. In these analyses, the mean FA value for the two animals was averaged across all diaphyseal regions (n = 12) and compared with the corresponding value for all metaphyseal regions (n = 12) or axial bones (n = 6) (see Table 1, note 2–3 for a list of the relevant bones or bone regions).

**Table 2.**
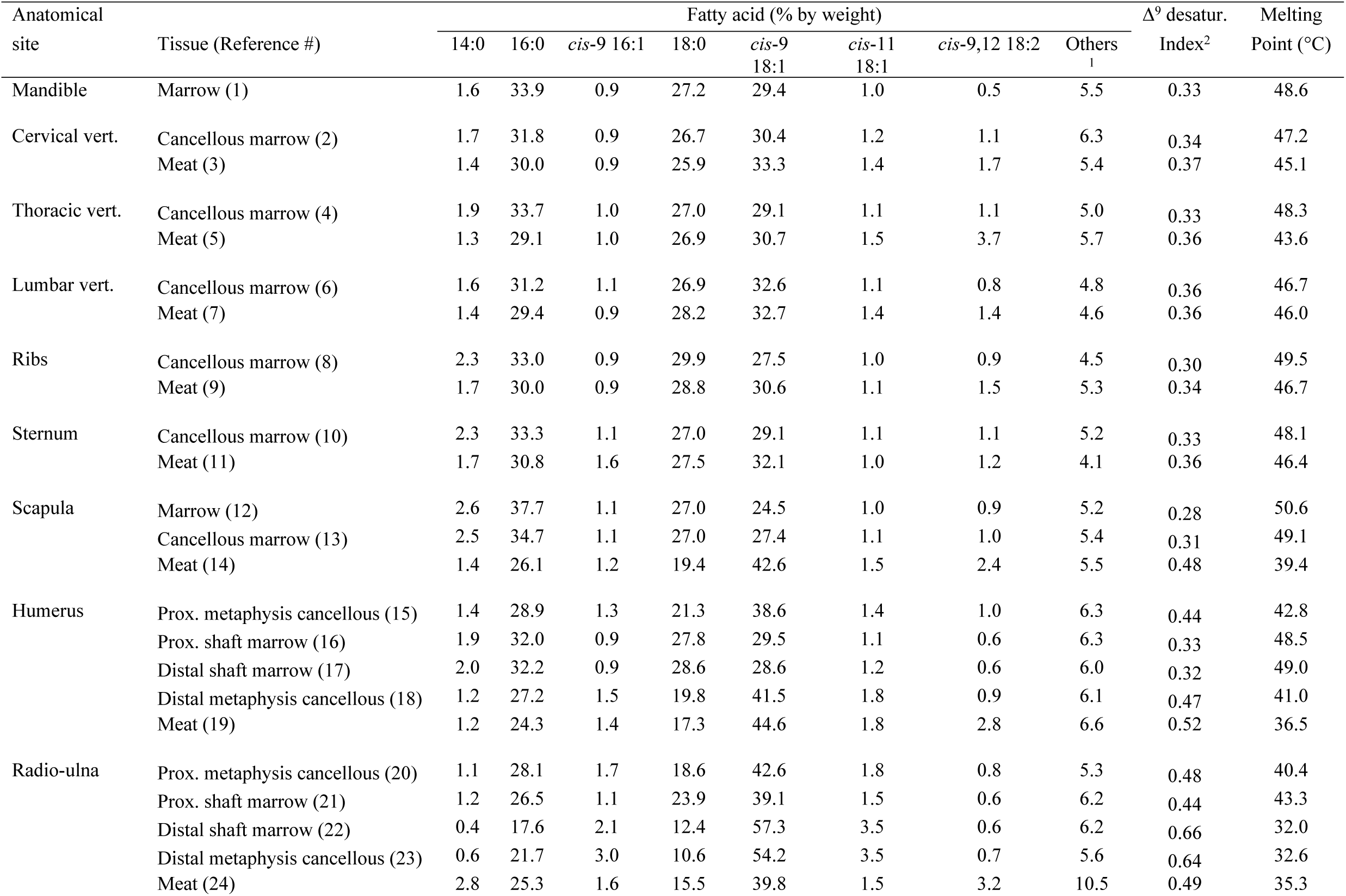
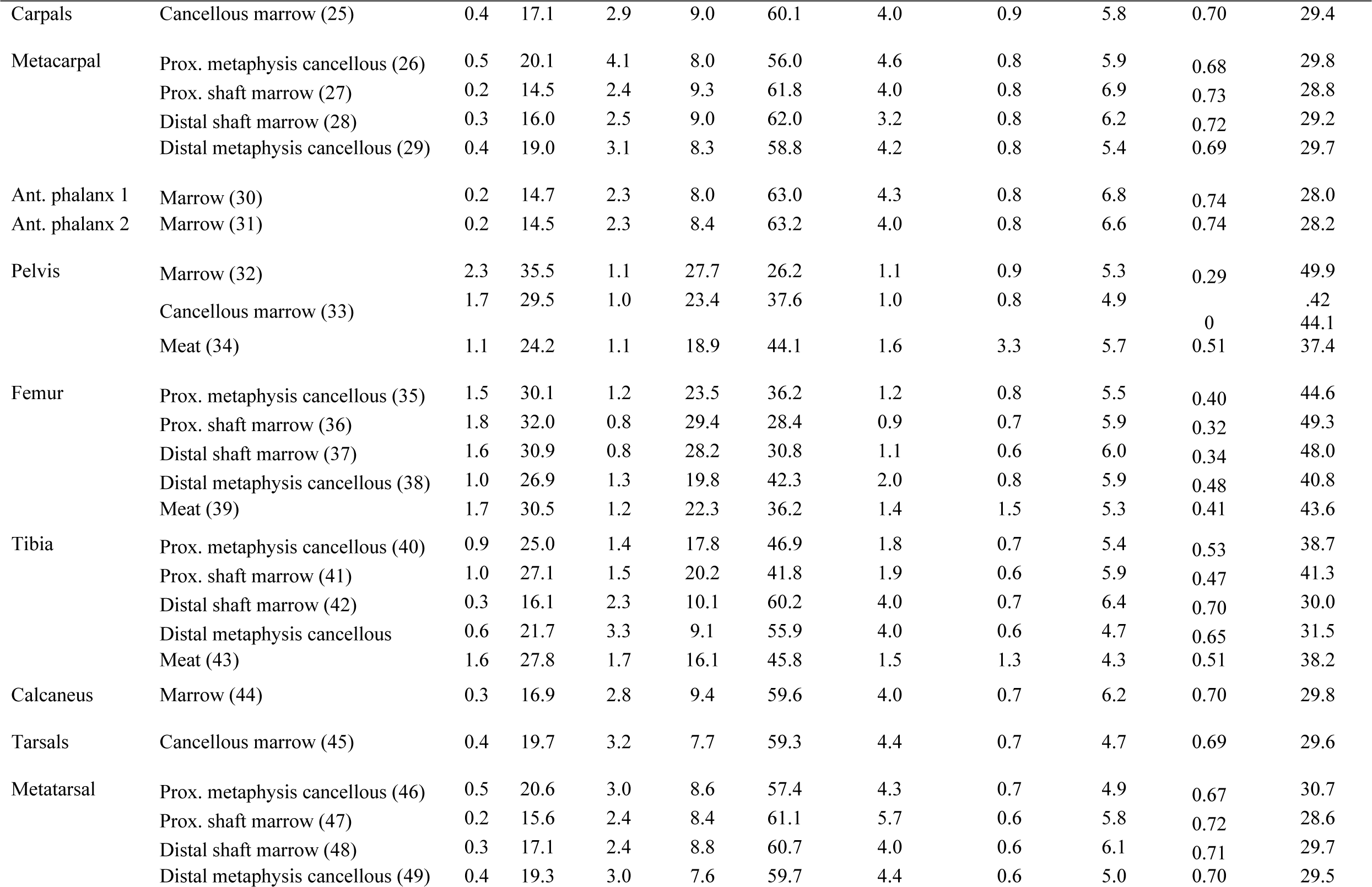
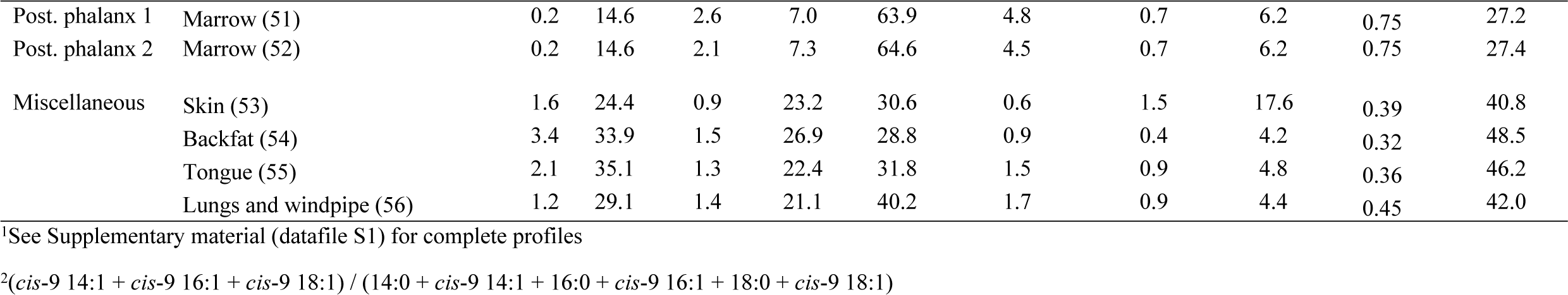
Fatty acid profile and calculated melting point of lipid extracted from varying body tissues in caribou. The values are averages for the two animals.

## Results

In the shaft cavity of the long bones, the percentage of total lipids is high (74.9 ± 5.0%, n = 12, Table 1) as is the percentage of FA calculated on a crude fat basis (80.2 ± 4.5%, n = 12, Table 1). In comparison, the percentage of FA is significantly lower in the metaphyseal portions of the same bones (73.8 ± 4.8%, n = 12, Table 1, t = 3.37, *p* = 0.0028) and the axial skeleton (74.6 ± 4.3%, n = 6, Table 1, t = 2.523, *p* = 0.0226). These lower values are likely due to an increased representation of membrane lipids in the latter samples, which is consistent with their known hematopoietic function. However, the metaphyseal regions of the distal metapodials (metatarsals and metacarpals) may represent an exception to this trend as they show only minor differences in FA composition when compared to the adjacent diaphyseal marrow (Fig 2a). We also note that the muscle tissues show low percentages of total lipids and FA, which suggests a low proportion of adipocytes in these tissues.

**Fig 2.**
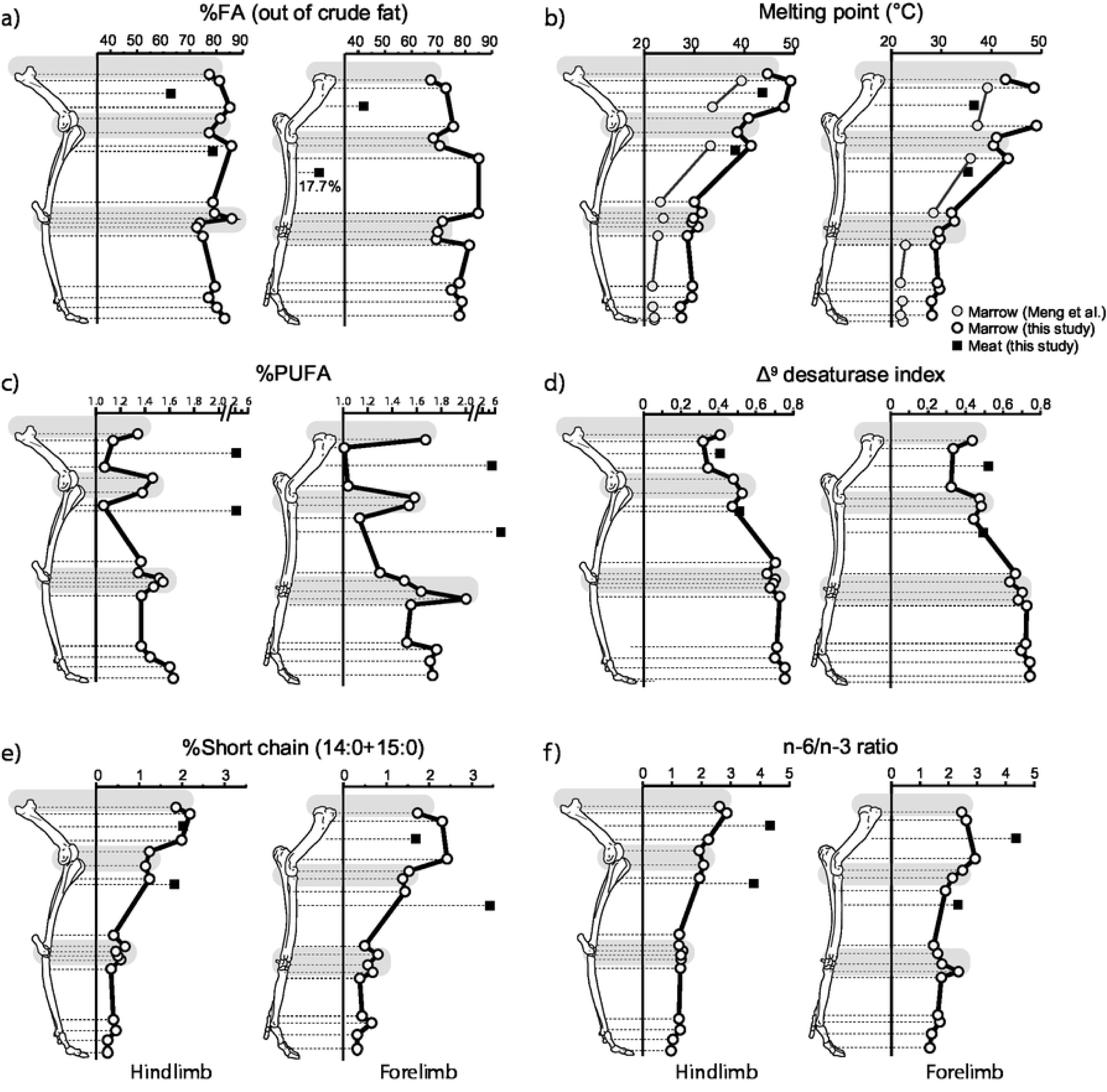
FA composition in the caribou limbs: a) percentage of FA (calculated out of crude fat); b) weighted melting point; c) percentage of polyunsaturated FA (PUFA); d) Δ^9^ desaturase index; e) %short chain saturated FA; f) n-6/n-3 FA ratio. Note the change of scale in c). The shaded areas correspond to areas of suspected significant hematopoiesis. In this and the following figure, the data points correspond to the mean for the two animals. The summed percentage of PUFA (%PUFA) includes these FA: *cis*-9,12 18:2; *cis*-9,12,15 18:3; *cis*-6,9,12,15 18:4; *cis*-11,14 20:2; *cis*-8,11,14 20:3; *cis*-5,8,11,14 20:4; *cis*-5,8,11,14,17 20:5; *cis*-7,10,13,16 22:4; *cis*-4,7,10,13,16 22:5; *cis*-7,10,13,16,19 22:5; *cis*-4,7,10,13,16,19 22:6. The percentage of short chain saturated FA (%short chain saturated FA) is the sum of 14:0 and 15:0. The Δ^9^ desaturase index was derived as follows: (*cis*-9 14:1 + *cis*-9 16:1 + *cis*-9 18:1)/(14:0 + *cis*-9 14:1 + 16:0 + *cis*-9 16:1 + 18:0 + *cis*-9 18:1). The n-6/n-3 ratio was obtained using this equation: n-6 PUFA/n-3 PUFA. Data from Table 2–3 and Supplemental material (datafile S1).

The gas chromatography analysis allowed the identification and quantification of 33 different FA varying from 14 to 22 carbon chain lengths (Table 2). In the appendicular skeleton, we note a gradual increase in the proportion of the *cis*-9 monounsaturated FA (MUFA) as one progresses away from the body core (Table 2). This increase is primarily expressed in the form of a greater representation of oleic acid (*cis*-9 18:1) and, to a lesser extent, palmitoleic acid (*cis*-9 16:1) in the extremities. Conversely, a gradual decrease in several classes of saturated FA— mostly palmitic acid (16:0) and stearic acid (18:0)—is observed distally. These changes in the FA composition of appendicular marrow produce a steady decrease in average fat melting point as one moves toward the extremities (Fig 2b). However, the average melting point of the metaphyseal regions is consistently lower than predicted by the FA pattern for diaphyseal marrow (Fig 2b, Tables 2–3). These lower melting points may signal an increased presence of lipids from membranes in metaphyseal regions, which is consistent with the higher percentages of PUFA observed in the same regions (Fig 2c). When compared to the shaft regions and excluding the metapodials, the metaphyseal regions also show higher values for the Δ^9^ desaturase index (Fig 2d) and lower percentages of short chain saturated FA (Fig 2e). In contrast, changes in the n-3/n-6 ratio are small (Fig 2f).

A comparison of the limbs with the axial skeleton shows several interesting patterns. In muscle tissues, values for the Δ^9^ desaturase index are systematically lower in the body core (tongue to sternum: 0.360 ± .011, n = 6) than in the limbs (scapula to tibia: 0.486 ± .041, n = 6, t = 7.2705, *p* < 0.0001, Fig 3a), a trend also observed in cancellous marrow (cervical to sternum: 0.332 ± .021, n = 5; scapula to distal metatarsal: 0.559 ± .132, n = 16, t = 3.7649, *p* = 0.0013, Fig 3b). We note that the percentage of short chain saturated FA (Fig 3c) and the n-6/n-3 ratio (Table 3) are higher in the cancellous marrow of the axial skeleton than that of the limbs where patterns of steady decrease are observed as one proceeds distally (Fig 3c–d).

**Fig 3.**
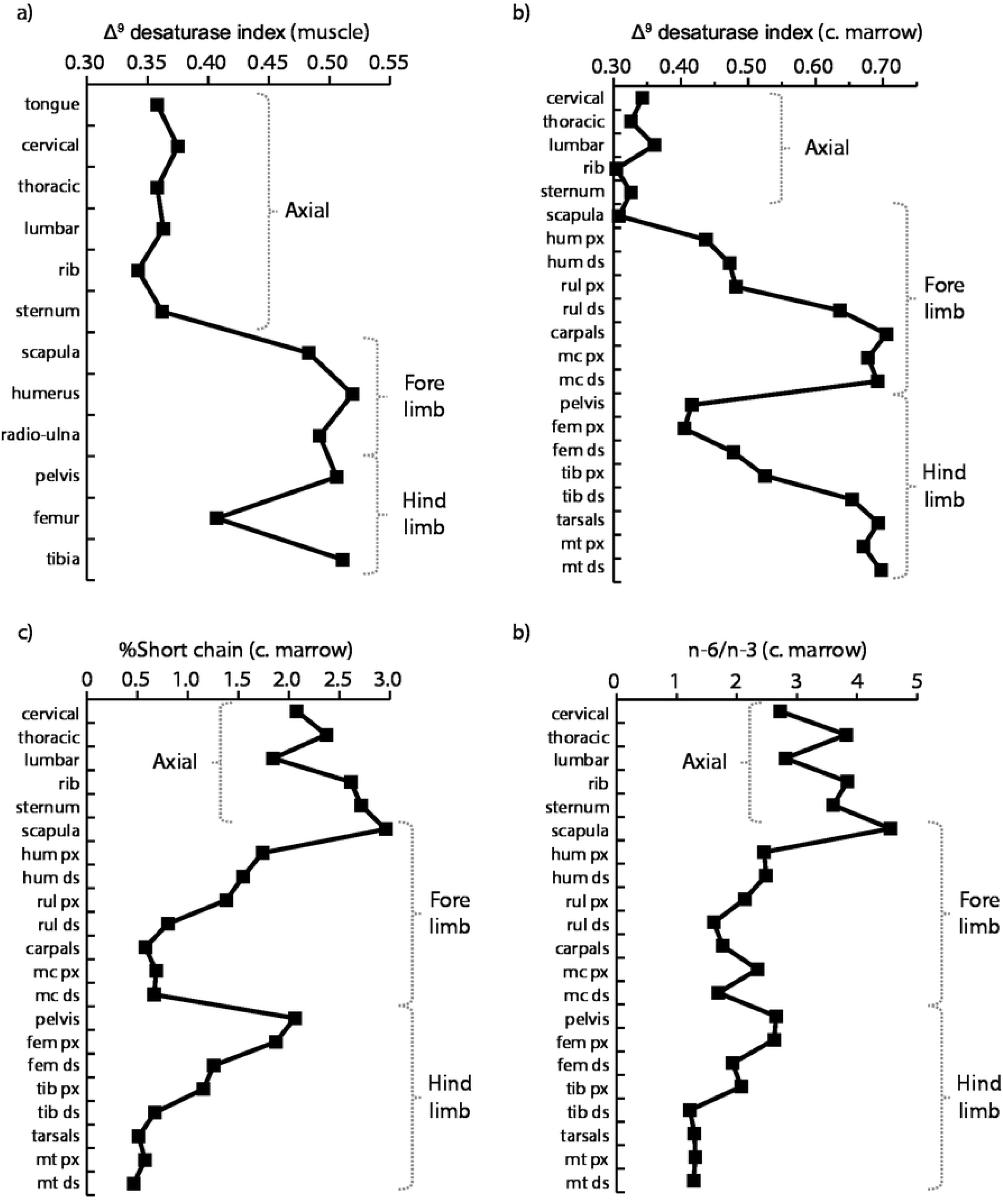
FA in the axial and appendicular skeleton: a) Δ^9^ desaturase index in muscle tissues; b) Δ^9^ desaturase index in cancellous marrow; c) percentage of short chain saturated FA in cancellous marrow; d) n-6/n-3 ratio in cancellous marrow. Comparisons for marrow exclude the shaft cavities. Data from Tables 2–3 and Supplemental material (datafile S1). Abbreviations: c. marrow=cancellous marrow; hum=humerus; rul=radio-ulna; mc=metacarpal; fem=femur; tib=tibia; mt=metatarsal; px=proximal metaphysis; ds=distal metaphysis.

**Table 3.**
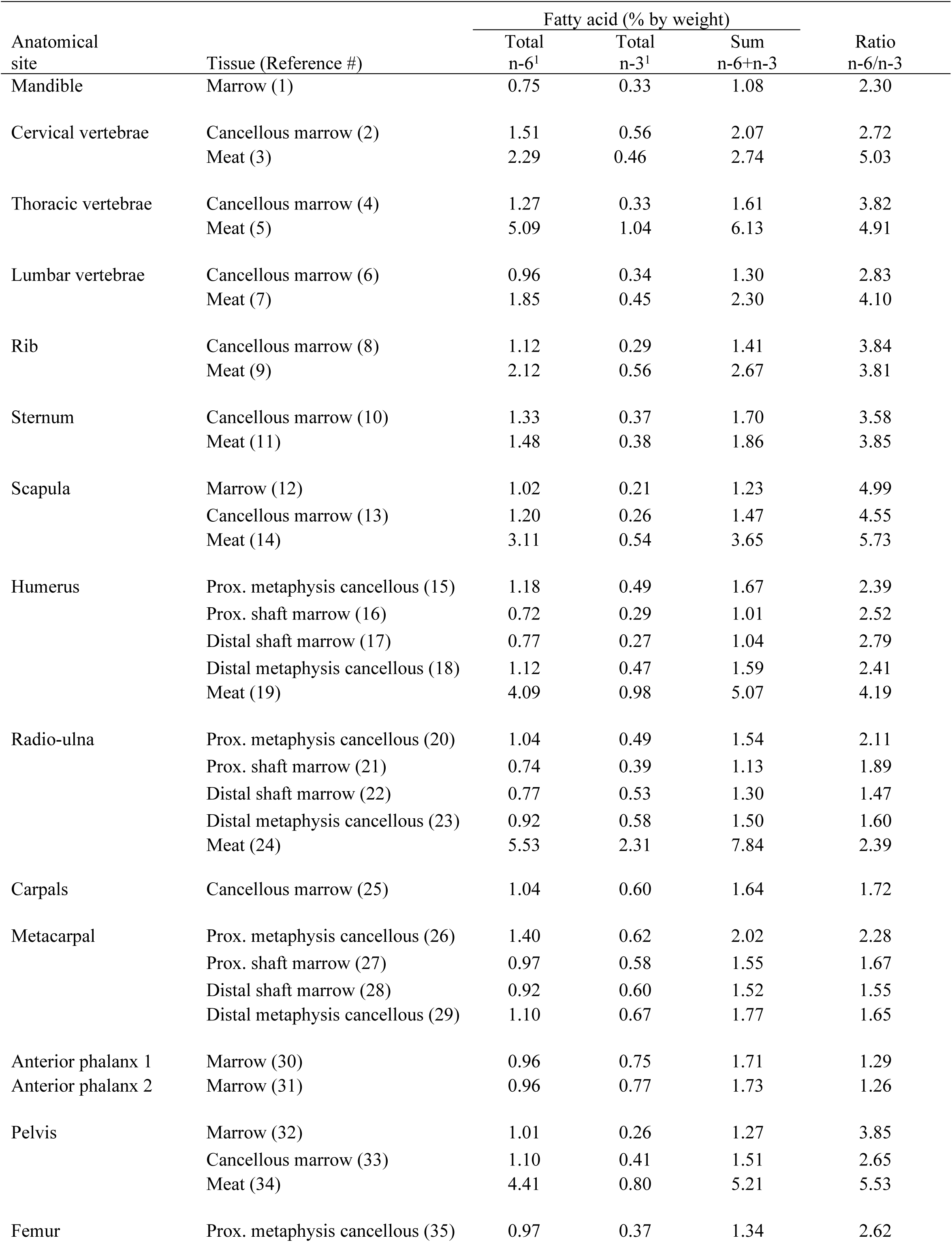
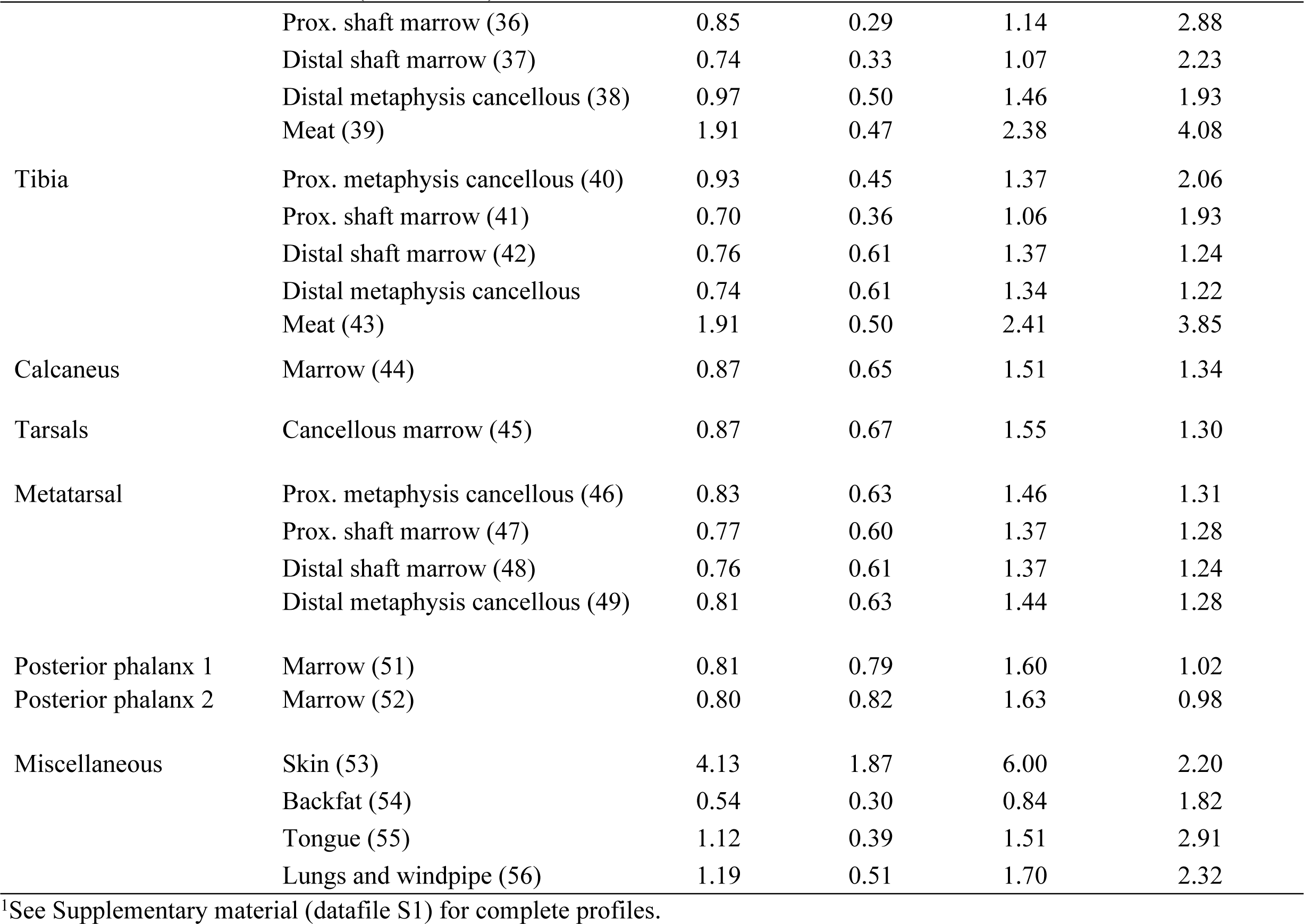
Polyunsaturated fatty acids family in lipids extracted from various body tissues in caribou.

## Discussion

Previous research on the thermal adaptation of limbs in cold-adapted mammals has largely focused on FA variation in the cBMA of long bones with occasional comparisons with other types of soft tissues (Meng et al. 1969; Käkelä and Hyvärinen 1996; Soppela and Nieminen 2001). In the present study, the FA profiles of a wide range of lipid-containing tissues in caribou were compared between and within different anatomical locations. Most of the bone marrow and muscle samples that we examined varied in terms of lipid proportions. The FA profiles suggest that the diaphyseal marrow and backfat are dominated by triacylglycerols derived from adipocytes (loosely connected with collagen) whereas the other tissues—such as muscles, and perhaps, rBMA—appear to show greater proportions of phospholipids and cholesterol esters derived from cell membranes.

In comparison to the shaft regions, the metaphyseal regions show higher values for the Δ^9^ desaturase index, higher percentages of PUFA and lower percentages of short chain saturated FA. These trends are compatible with significant hematopoietic activity in the articular ends of the long bones (Tavassoli and Yoffey 1985) and with increased desaturation in the extremities of caribou/reindeer (Meng et al. 1969; Pond et al. 1993; Soppela and Nieminen 2001). In agreement with Meng et al.’s (1969) observations about the FA composition of cBMA, the diaphyseal regions of the long bones show a pattern of desaturation toward the extremities, which contributes to lowering the melting point of the adipose tissues. A similar trend is seen in the metaphyseal regions. The distal decrease in melting point in diaphyseal and metaphyseal regions supports the hypothesis of a physiological adaptation of the lipid component of cells to the marked heterothermia that may be displayed in reindeer legs (Irving & Krogh 1955, Johnsen et al. 1985). This is because a low melting point allows for soft tissues of the peripheral parts— including extremities that are less insulated and are allowed to cool more than the trunk in order to limit heat loss rate—to remain supple when facing cool thermal conditions.

It is known that FA with a higher degree of unsaturation—especially those high in PUFA with their double bonds located near the methyl end—can, when need arises, be mobilized more readily than saturated FA (Gavino and Gavino 1992; Raclot and Groscolas 1993; Connor et al. 1996; Raclot et al. 1995). For instance, in situations of impaired energy balance, unsaturated and long-chain FA are preferentially mobilized from triacylglycerols of adipose tissues (Soppela and Nieminen 2002, Nieminen et al. 2006, Mustonen et al. 2009), including bone marrow fat (Soppela and Nieminen 2001) and brown adipose tissue (Groscolas and Herzberg 1997). However, additional work will be needed to assess whether there are differences in the fat mobilization process between rBMA and cBMA.

Our analysis also shows important differences between the metaphyseal and diaphyseal portions of the long bones, the former regions showing higher percentages of PUFA, higher values for the Δ^9^ desaturase index and lower melting points. In the caribou females that we sampled, both scored as prime adults, the metaphyseal regions of the long bones were apparently actively involved in hematopoiesis. If confirmed, this result would be consistent with observations made in humans and many other mammals (Tavassoli and Yoffey 1985).

Previous studies that compared different muscles in terrestrial mammals have often stressed the lack of FA variation in intramuscular fat between muscles of single animals (Nikolaidis and Mougios 2004; Wood et al. 2008) and in patterns of FA desaturation between species living at different latitudes (Guerrero and Rogers 2019). Our results show relatively minor changes in FA composition between different muscles in the axial skeleton. However, the Δ^9^ desaturase index shows clear and apparently systematic differences in the muscles between the axial and appendicular skeleton. The pattern that we uncovered is consistent with appendicular muscle tissues being more unsaturated than those of the axial skeleton (Pond et al. 1992, 1993; Mustonen et al. 2007), presumably as an adaptation to the cooler temperature seen in the limbs of mammals due to their thermoregulatory vasoconstrictor responses aimed at minimizing limb heat loss. As a last point, it is well established that PUFA are critical in influencing the fluidity, permeability and protein binding functions of cell membranes. The changes in PUFA abundance seen in the bone marrow and muscle tissues that we examined are in agreement with this interpretation.

## Conclusion

It is increasingly clear that the rBMA in the metaphyseal regions differ morphologically and functionally from that found in the cBMA of diaphyseal regions (Scheller et al. 2015; Craft et al. 2018). Assuming that our interpretation of BMA distribution in the limb samples is correct, our analysis suggests that cBMA and rBMA vary systematically in terms of FA composition, with both subtypes of adipocytes showing a distal increase in the degree of unsaturation and an overall decrease in fat melting point in the limbs. While variation in FA composition seems limited in muscles of the body core, the cell membranes of muscle tissues show patterns of change of FA in the limbs, including an increase in the Δ^9^ desaturase index, that are consistent with an adaptation to exposure to ambient temperature. Given that patterns of thermoregulation seem widely shared among terrestrial mammals—with frequent contrasts being seen between the warmer core and the more heterothermic extremities and appendages—we suggest that the trends of FA composition that we observed in the appendicular skeleton of caribou is also characteristic of other species, possibly including humans.

## Acknowledgments

Funding was provided through a grant from the Laboratoire d’Archéométrie de l’Université Laval. We would like to thank Lars Folkow, Sanna-Mari Kynkäänniemi, Vassili Mougios, Petteri Nieminen, Cara Ocobock, Jürgen Steinmeyer and Glenn Tattersall for their insightful observations and several useful discussions regarding the implications of our data.

## List of Supplements

Datafile S1. Complete FA profiles for the two caribou individuals.

